# Plasma derived cell-free mitochondrial DNA originates mainly from circulating cell-free mitochondria

**DOI:** 10.1101/2021.09.03.458846

**Authors:** Benoit Roch, Ekaterina Pisareva, Cynthia Sanchez, Brice Pastor, Rita Tanos, Alexia Mirandola, Thibault Mazard, Zahra Al Amir Dache, Alain R. Thierry

## Abstract

Circulating mitochondrial DNA (cir-mtDNA) could have a potential comparable to circulating nuclear DNA (cir-nDNA), with numerous applications. However, research and development in this area have fallen behind, particularly considering its origin and structural features. To tackle this, we initially combined Q-PCR and low-pass whole genome sequencing in the same analytical strategy previously and successfully used for cir-nDNA. This revealed unexplained structural patterns and led us to correlate these data with observations made during physical examinations such as filtration, and differential centrifugation in various plasma preparations. Both the integrity index and number of reads revealed a very minor proportion of low size-ranged fragments (<1000 bp) in plasma obtained with a standard preparation (0.06%). Filtration and high speed second step centrifugation revealed that 98.7 and 99.4% corresponded to extracellular mitochondria either free or in large extracellular vesicles. When avoiding platelet activation during plasma preparation, the proportion of both types of entities was still preponderant (76-80%), but the amount of detected mitochondrial DNA decreased 67-fold. In correlation with our previous study on the presence of circulating cell-free mitochondria in blood, our differential centrifugation procedure suggested that cir-mtDNA is also associated with approximately 18% small extracellular vesicles, 1.7% exosomes and 4% protein complexes.

## INTRODUCTION

Circulating DNA (cirDNA) possesses considerable potential for the study of both healthy subjects and patients with underlying pathological conditions (1–3). Thus, recent adv5nces in the understanding of cirDNA have seen its use broadened, leading to the design of numerous specific approaches, as is testified by its numerous current clinical applications (4–9). CirDNA has been intensely studied, with significant efforts being made to improve its detection and to discriminate its tissue/cells of origin, so that its diagnostic potential may be optimized (8,10,11). CirDNA derives from both nuclear DNA (nDNA) and extrachromosomal mitochondrial DNA (mtDNA) (10–13). However, most studies were restricted themselves mainly to nuclear cirDNA (cir-nDNA), as compared to mitochondrial cirDNA (cir-mtDNA).

Mitochondria, often referred to as the powerhouses of the cell, help turn the energy coming from the food intake into energy usable by the cell. Mitochondria are also involved in signaling between cells, cell death, heat production, and signaling calcium (14,15). Given its multiple physiological functions, mitochondria has been considered as a potential biomarker of many normal and pathological conditions (14,16,17). Beside full mitochondria detection, most studies towards this goal have focused on mitochondria constituents, principally on the mtDNA content. Cellular mtDNA is accessible from whole blood or peripheral blood mononuclear cells (PBMCs), and has been quantified across multiple clinical areas (13,14,16,18,19). Consequently, with the growing interest in cirDNA research and applications, cir-mtDNA is now positioned as a potentially strong biological source in the diagnostic field (20).

What is true for cir-nDNA, regarding improved knowledge of its structures leading to improved detection, should also be true for cir-mtDNA. Investigation on cirDNA structures has focused mainly on cirDNA fragmentation (fragmentomics), since elucidation of the cirDNA fragment size distribution may reveal characteristics linked to their release mechanism, as well as the protection against degradation in the blood stream, which is provided by DNA packaging in nucleo-proteic/lipidic complexes. While DNA is highly sensitive to DNase in a biological environment (21,22), its highly negatively charged molecules have a significant capacity to bind, reversely condense, and pack tightly into macromolecular structures. nDNA is packed within nucleosomes and condensed in a hierarchical and tunable architecture mediated by DNA-protein interaction constituting the chromatin in Archaea and eukaryotes (23). Holdenrieder et al. suggested that cirDNA were mainly packed in nucleosomes (24), and several reports have shown that cirDNA associated structures have nucleosome footprints (25–27). We recently demonstrated that cirDNA are mainly compacted within mono-nucleosomes, which apparently constitutes their most stable form, while di- or oligo-nucleosomes or larger pieces of cirDNA constitute a very minor fraction of its population (28,29). In addition, we demonstrated that blunted and jagged double-stranded DNA (dsDNA) of size up to 220 bp and down to 70 bp are packed in nucleosome/chromatosome particles (29). Q-PCR assisted data on cirDNA distribution corresponded to data obtained from low-pass whole genome sequencing (WGS) performed using single-stranded DNA (ssDNA) library preparation (SSP) rather than dsDNA library preparation (DSP), allowing the harmonization of data obtained from both techniques (28).

Despite all of this, we still understand far less about the characteristics of cir-mtDNA than those of cir-nDNA, especially its topology in circulation. In a recent study performed by our team, we showed that the mitochondrial genome may be found in nearly 50,000-fold more copies than the nuclear genome in the plasma of healthy individuals (30). This suggests the existence of stabilizing structures protecting mtDNA molecules, thus allowing the detection and quantification of cir-mtDNA in the bloodstream (10,16,31,32). We later demonstrated that blood contains cell-free intact mitochondria as well as cir-mtDNA (30). Due to the lack of histone in mitochondria, there is so far no full explanation for cir-mtDNA stabilization/protection in the blood circulation.

The accurate differentiation of cirDNA of nuclear and mitochondrial origin is feasible (10,31,33), and may offer diagnostic information in specific physiological or pathological situations (32,34,35). Note, we hypothesized that the respective quantitation of cir-mtDNA and cir-nDNA may have potential cancer screening capacity (10,36,37). Consequently, an elucidation of the structural features of cir-mtDNA may improve their detection and quantification. Thus, a major focus of our work is to determine if the profile of detected cir-mtDNA really matches one previously described as a trash profile (26,38), and also to determine if this profile is reproducible between subjects.

To this end, we first explored cir-mtDNA’s structural features, using the analytical strategy of combined Q-PCR and low-pass WGS (LP-WGS) analysis, from both DSP and SSP, which provides a determination of the fractional distribution over a wide size range (28,29). This pointed to unexplained structural features, and led us to correlate these data with observations made using physical examination. This enabled us to decipher the forms and structures with which mtDNA is associated in blood circulation. In doing so, our work revealed profound differences between cir-mtDNA and cir-nDNA in terms of size distribution, structure and origin. This led us to significantly revise previous assumptions regarding the signification/characterization of cir-mtDNA, which was previously used as a potential biomarker of numerous pathologies (33,35,39,40).

## MATERIALS AND METHODS

### Sources of blood samples

The French Blood Establishment (EFS) of Montpellier provided blood draws from healthy individuals (HI). An informed consent form was signed for all individuals.

### Preanalytical work-up

By definition plasma is the liquid that remains when blood clotting is prevented by the addition of an anticoagulant. Pre-analytical conditions used to obtain a plasma fraction in this way can differ, although all include a centrifugation step to remove cells. Various processes were used in the study.

#### Standardized plasma preparation (SPP) for cirDNA analysis (cirDNA-SPP)

A standard protocol for plasma isolation in EDTA tube by double centrifugation was previously used in numerous studies, following initial validation by Chiu et al (41) and El Messaoudi et al (42). Briefly, samples were collected in EDTA tubes and handled while respecting stringently pre-analytical guidelines previously established (43). Blood samples were consecutively submitted to a double centrifugation at 4°C for 10 minutes: first at 1,200 g (low speed centrifugation, LS) to obtain a plasma that then underwent a second 16,000 g centrifugation (high speed centrifugation, HS). Subsequent supernatant was carefully removed down to 1 cm from the cell layer (43). CirDNA was then extracted from 200μL of this plasma preparation in a final volume of elution of 80μL, using QIAamp DNA Blood Mini Kit (Qiagen). DNA extracts were immediately stored at −20°C until use. Filtration (F) with 0.22μM Polysulfone membrane filter (30) and a freezing step at −20C° were also performed between the two centrifugation steps to study cirDNA origins.

#### Plasma preparation without platelets activation (PPw/oPA)

Fresh blood was collected in BD Vacutainer citrate-theophylline-adenosine-dipyridamole (CTAD) tubes (Ozyme, Montigny-le-Bretonneux, France). Plasma was isolated via differential centrifugations, all performed for 10min at room temperature (RT) without interruption: two successive centrifugations were first performed at 200 g, followed by a third centrifugation at 300 g. Preheated (37°C) Anticoagulant Citrate Dextrose Solution A (ACD-A) buffer (0.1M trisodium citrate, 0.11M glucose and 0.08M citric acid) and Prostaglandin E1 (1μM) (Sigma-Aldrich, St. Quentin Fallavier Cedex) were then added to the plasma, which was then further centrifuged at 1,100 g, and finally centrifuged again at 2,500 g. CirDNA analysis was performed on the plasma preparation obtained from the supernatant following an additional centrifugation step at 16,000 g for 10min at RT.

#### Differential centrifugation and filtration of plasma

Various plasma preparations from a single blood draw for each individual were tested in respect to centrifugation speed and filtration. First, the plasma isolated following the 1,200 g centrifugation as above described was aliquoted in 4 equal volumes: (i), one aliquot was considered as a control (LS) with no additional treatment; (ii) one was filtered (LS+F) with a 0.22μM filter (Sartorius Minisart High Flow, Fisher Scientific, Illkirch, France); (iii), the third one was centrifuged at 16,000 g for 10min at 4°C (LS+HS); and (iv), the fourth one was stored at −20°C before being subjected to the 16,000 g centrifugation (LS+freezing+HS). The cirDNA was extracted from previous plasma preparations and then analyzed by Q-PCR. Second, the plasma was isolated at 400 g with Ficoll gradient. The supernatant was centrifuged at 16,000 g for 10min at 4°C, (Micro Star microcentrifuge, VWR), then further centrifuged at 40,000 g for 1h at 4°C, and finally centrifuged at 200,000 g for 2h at 4°C (Beckman MLA-130 Ultracentrifuge Rotor). After each centrifugation step, an aliquot was performed and the cirDNA extracted from supernatants for Q-PCR analysis.

### Quantification of cir-mtDNA and cir-nDNA by Q-PCR

Q-PCR amplifications were performed, at least in duplicate, on a CFX96 Touch Real-Time PCR Detection System instrument using the CFX Manager Software 3.0 (Bio-Rad). The reaction volume was 25μL with each PCR reaction mixture composed of 12.5μL PCR mix (Bio-Rad SsoAdvanced Universal SYBR Green Supermix), 2.5μL of each amplification primer (0.3pmol/μL, final concentration), 2.5μL PCR-analyzed Nuclease Free Water (Qiagen), and 5μL of DNA extract.

Thermal cycling started with a hot-start polymerase activation denaturation step performed at 95°C for 3min, followed by 40 repeated cycles with a succession of 90°C for 10s and 60°C for 30s. An incremental increase of the temperature from 55°C to 90°C, with a plate reading every 0.2°C, produced the melting curves. To calibrate the quantifications, we used serial dilutions of genomic DNA from the DIFI human colorectal cancer cell line at known concentrations, for each pair of primers. The genomic DIFI cell line was used as a reference to assess the efficiency of these primers. For each targeted size DNA, the concentrations measured correspond to the number of amplicons obtained in each well. Sample concentrations were determined from triplicate measurements and extrapolated from these standard curves. A negative control was set in each run for each pair of primers.

Concentrations of cir-nDNA and cir-mtDNA were determined in seven HI using a nested Q-PCR primer system to detect amplicons, respectively targeting validated wild type sequences in the *KRAS* gene (67 bp and 320 bp) (44,45) and in the *MT-CO3* gene (67 bp and 310 bp) (10,37) (Table 1). Calibration of mitochondrial and nuclear genome equivalent copy numbers were determined as previously described (10). Q-PCR experiments followed MIQE guidelines (46).

**Table 1:** Circulating DNA (cirDNA) mean concentrations and size markers in seven healthy individuals (ID 1-7), as determined in plasma by Q-PCR. Mean concentrations of nuclear (cir-nDNA) and mitochondrial cirDNA (cir-mtDNA), expressed in ng/mL, were determined using a nested Q-PCR primer system to detect amplicons of different sizes, targeting *KRAS* gene and *MT-CO3* gene, respectively. For cir-mtDNA, amplicons of 67 bp (short) and 310 bp (long) targeted *MT-CO3* gene. For cir-nDNA, amplicons of 67 bp (short) and 320 bp (long) targeted a *KRAS* intron 3/exon 2 sequence. Mitochondrial DII (mtDII) and nuclear DNA Integrity Index (nDII) represent the respective ratio of mean plasma concentrations of long over short amplicons.

| ID sample | cir-mtDNA (ng/mL)<br>plasma | cir-nDNA (ng/mL)<br>plasma | Mean cir-mtDNA (%) from total | mtDII | nDII |
| --- | --- | --- | --- | --- | --- |
| 1 | 0.0044 | 1.37 | 0.32 | 1.05 | 0.48 |
| 2 | 0.0014 | 1.51 | 0.09 | 0.99 | 0.34 |
| 3 | 0.0024 | 1.50 | 0.16 | 0.98 | 0.22 |
| 4 | 0.0033 | 0.57 | 0.58 | 0.96 | 0.34 |
| 5 | 0.0060 | 1.34 | 0.44 | 1.07 | 0.10 |
| 6 | 0.0070 | 4.60 | 0.15 | 1.03 | 0.09 |
| 7 | 0.0080 | 4.95 | 0.16 | N/A | 0.16 |
| Mean | 0.0046 | 2.26 | 0.27 | 1.01 | 0.25 |
| SD | 0.0025 | 1.75 | 0.18 | 0.04 | 0.14 |

### DNA integrity Index (DII)

The DII was determined using aforementioned nested Q-PCR primer systems. The overall cirDNA fragmentation level estimated by the DII was previously validated for cir-nDNA (45,47–49) and for cir-mtDNA (10,36,37). DII is based on the ratio of the concentration of the longest fragments, respectively 310 bp for cir-mtDNA and 320 bp for cir-nDNA, over this of the shortest fragments, 67 bp in both cir-mtDNA and cir-nDNA, in each respective DNA region studied. It corresponds to the fraction of cirDNA fragments whose size is over the longest amplicon.

### Preparation of sequencing libraries

Both DSP and SSP were prepared. SSP allows the integration of ssDNA and dsDNA in the library. DSP libraries were prepared with the NEB Next® Ultra™ II kit (New England Biolabs). SSP libraries were prepared with the Swift ACCEL-NGS® 1S PLUS kit (Swift Biosciences). For both preparations, a minimum of 1ng of cirDNA was engaged without fragmentation, and each kit providers’ recommendations were followed. For DSP, repaired A-tailed fragments were submitted to a ligation with Illumina paired-end adaptor oligonucleotides. A purification was then applied by solid-phase reversible immobilization (SPRI), followed by an enrichment by 11 PCR cycles with unique dual index (UDI) primers indexing, and another SPRI purification. For SSP, to convert all DNA into single strands, a first step of heat denaturation was accomplished. Through the use of an adaptase, this protocol enables the simultaneous binding of an adapter to the end of each single-stranded fragment and the lengthening of the 3’ end of this fragment. Subsequently, a primer extension synthesized the complementary strand, followed by a SPRI purification, the ligation of a second adapter at the other end, and another step of SPRI purification. An enrichment of the product was then performed by 11 PCR cycles, followed by a final SPRI purification. An adjustment of the SPRI purification was operated to keep the small fragments around 70 bp of insert for both types of preparation. Lastly, a precise quantification by Q-PCR was applied to the libraries to be sequenced, to ensure that the appropriate DNA quantity was loaded to the Illumina sequencer, and that a minimum of 1.5 million of clusters would be obtained.

The calculation of each fragment size’s frequency, expressed in %, was carried out using the ratio of the sequenced reads to the total reads obtained in the library. The fragment size distribution length unit was base pairs (bp) when using DSP-sequencing (DSP-S), and nucleotides (nt) when using SSP-sequencing (SSP-S). To allow the comparison of data from both DSP-S and SSP-S, the size or size range were then expressed as bp (nt).

Note, these are PCR-free libraries, with no initial PCR targeted sequencing performed, given our knowledge on the previously described high cirDNA fragmentation showing an appropriate mean fragment size range for WGS, from 150 to 160 bp (28).

### Size profile analysis by low-pass WGS

A NextSeq 500 Sequencing System or NovaSeq 6000 Sequencing System (Illumina) were used to sequence all libraries as paired-end 100 reads. Image analysis and base calling was achieved through the use of Illumina Real-Time Analysis, using default parameters. A cut of the individual barcoded paired-end reads (PE) was performed with Cutadapt v1.10, removing the adapters to discard trimmed reads shorter than 20 bp. An alignment of the trimmed FASTQ files with the human reference genome (Genome Reference Consortium Human Build 38, https://www.ncbi.nlm.nih.gov/grc/human) was obtained using the Maximum Entropy Method (MEM) algorithm in the Burrows-Wheeler Aligner (BWA) v0.7.15. The insert sizes were then extracted from the aligned bam files with the Template Length (TLEN) column for all pairs of reads with an insert size between 0 and 1000 bp.

The ratio of the number of reads at each fragment size to the total number of fragments from 30 to 1000 bp (nt) was used to calculate the frequency of each fragment size separated by one bp or nt. Relying respectively on the presence of dsDNA or ssDNA, the size profiles generated by DSP-S or SSP-S were respectively expressed in bp and nt as a fragment size unit. Note, the fragment sizes are offset by 3 bp (nt) for the two methods, which is consistent with damaged or non-flush input molecules, whose true endpoints are more faithfully represented in single-stranded libraries.

DNA libraries and sequencing were performed by IntegraGen SA (Evry, France). Under the aforementioned technical conditions, the limits of detection by sequencing are estimated to be 30 bp (nt) at the lower end, and nearly 1000 bp (nt) at the upper end. Sequencing of plasma-extracted cirDNA was performed from either DSP or SSP in plasma as previously reported (29).

### Statistics and drawings / Statistical analysis

Statistical calculation, log transformation of appropriate data and analysis were performed using GraphPad Prism software V6.01. The Student’s t-test was used to compare means. Differences with a two-sided p-value under 0.05 were considered to be statistically significant. In figures, significant p-values appear this way: *p<0.05, **p<0.01; ***p< 0.001; ****p< 0.0001. Figures of cirDNA size profile as performed by LP-WGS were drawn using R studio.

## RESULTS

### CirDNA concentration and size distribution in healthy individuals, as determined by Q-PCR, from plasma prepared with the cirDNA standard protocol

Cir-mtDNA and cir-nDNA total concentrations in plasma as determined by the short amplicon Q-PCR system averaged 0.0046 +/− 0.0025 and 2.26 +/− 1.75 ng/mL of plasma, respectively (Table 1). Mean concentration of cir-mtDNA from total cirDNA ranged between 0.09 and 0.58%, averaging 0.27 +/− 0.18%. Mitochondrial DII (mtDII) values were all nearly 1 (mean +/− SD 1.01 +/− 0.04), while nuclear DII (nDII) values were lower among HI plasma (mean +/− SD: 0.25 +/− 0.14) (Table 1).

### Low-pass WGS-based size profile of cirDNA, from plasma prepared with the cirDNA standard protocol

In HI, the mean number of reads for cir-mtDNA was 79 (range 24 – 207) for DSP-S and 119 (range 25-277) for SSP-S; the mean number of reads for cir-nDNA was 1,434,487 (range 1,079,717-1,611,205) for DSP-S and 1,007,070 (range 708,192-1,299,291) for SSP-S (Suppl. Table 1). The DSP-S and SSP-S ratios of the mean number of cir-mtDNA reads over cir-nDNA reads were 0.006% and 0.012% in HI (Suppl. Table 1). Because of the low read number of cir-mtDNA, we presented size profiles with histogram values at each fragment size up to 1000 bp (nt).

#### Mitochondrial circulating DNA size profile

The seven HI size profiles obtained by DSP-S and SSP-S are pasted over each other in Figure 1A and B, respectively. Using DSP-S, relative frequencies distributed from 55 to 425 bp (Fig. 1A), with a sharp increase up to 120 bp, a subsequent progressive decrease with no fragments above 425 bp, and a slightly higher number of fragments from 90 to 260 bp. Using SSP-S, relative frequencies distributed from 50 to 475 nt (Fig. 1B), with a sharp increase up to 80 nt, followed by a progressive decrease in the number of fragments. Two distinct populations appeared in the profiles obtained by SSP-S: one monomodal population from 50 to 150 nt, and a population smear from 150 to 475 nt, with no fragments above 475 nt (Fig. 1B). Cir-mtDNA size profiles appeared to be homogeneous among the seven HI, both for DSP-S and SSP-S (Fig. 1A, B). The majority of detected fragments in HI DSP-S mean size profiles distributed from 90 to 260 bp (77.9% of total fragments), with the number of fragments sharply increasing from 90 to 120 bp, peaking at 120 bp, and decreasing slowly in the range beyond that (Fig. 1C). The majority of detected fragments in HI SSP-S mean size profiles distributed from 50 to 150 nt (74.4% of total fragments), with the number of fragments sharply increasing from 50 to 80 nt, peaking at 80 nt, sharply decreasing from 80 to 150 nt, then decreasing slowly beyond that (Fig. 1D). Differences emerged when the DSP-S and SSP-S mean size profiles were compared (Fig. 1E and F). First, DSP-S mean size profiles distributed on a slightly shorter range (90 – 425 bp) than SSP-S (50 – 475 nt). Maximal values as determined by both methods ranged from about 100 to 150 bp for DSP-S and from 70 to 100 nt for SSP-S. Thus, SSP-S detected a higher number of low-sized sequences with a high mean frequency of fragments between 50 and 120 nt (61.0% of total fragments), while DSP-S showed a low number of fragments in this range (16.7% of total fragments).

**Figure 1:**
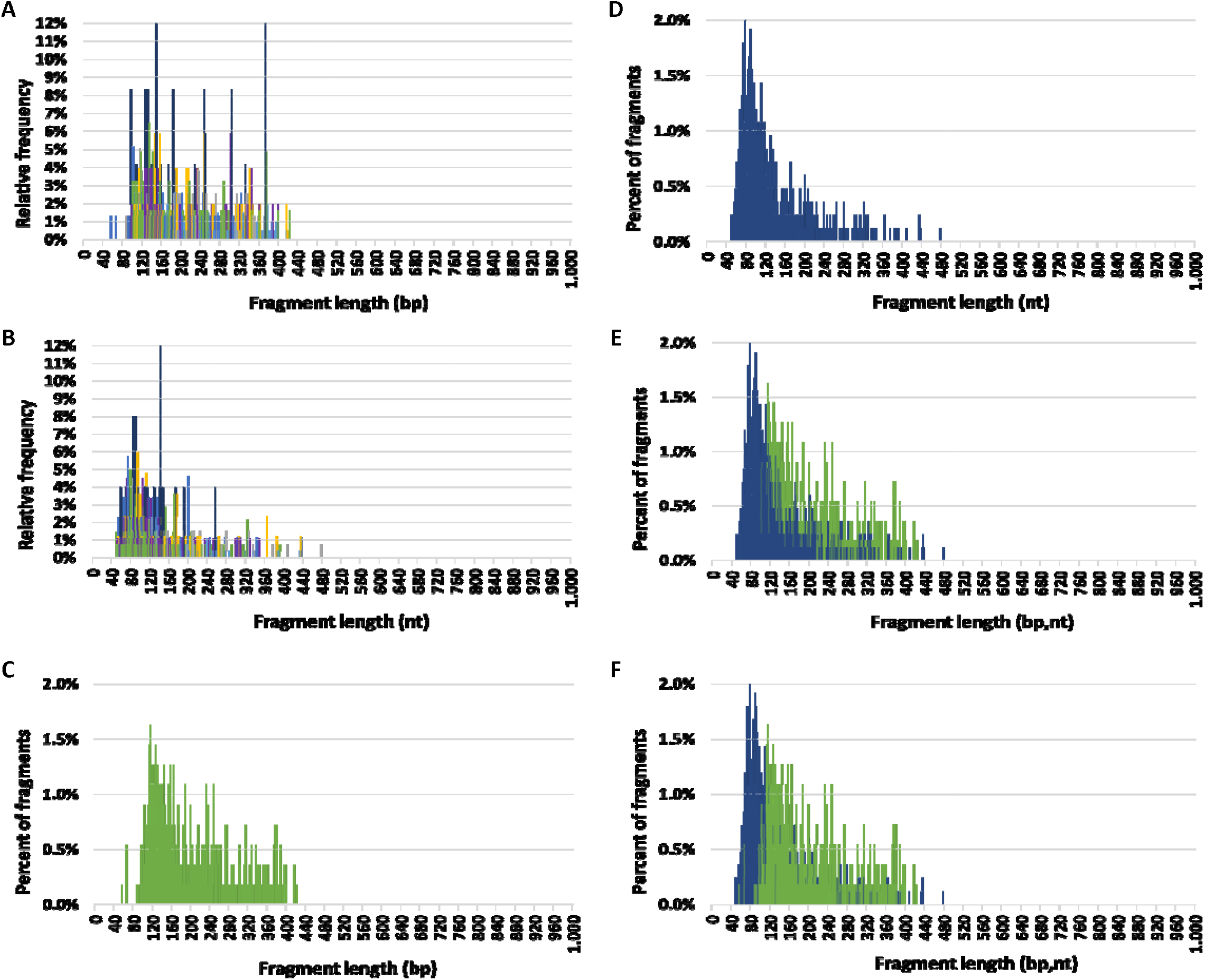
Circulating mitochondrial DNA (cir-mtDNA) size profile in healthy individuals. Relative frequencies of cir-mtDNA fragments from seven subjects (seven colours) as determined by low-pass WGS: A, DSP sequencing (DSP-S); B, SSP sequencing (SSP-S). Mean percent of fragments of cir-mtDNA from the seven subjects as determined by DSP-S (green, C, E, F) and SSP-S (blue, D, E, F): DSP-S alone (C), SSP-S alone (D), SSP-S in foreground and DSP-S in background (E), DSP-S in foreground and SSP-S in background (F).

#### Comparison of mitochondrial and nuclear circulating DNA size profiles

Cir-mtDNA and cir-nDNA DSP-S mean size profiles of HI ranged from 55 to 425 bp and 85 to 420 bp, respectively (Fig. 2A, B). The DSP-S cir-mtDNA profile of HI exhibited a major monomodal population, mostly ranging between 90 and 260 bp, peaking at 120 bp, and a population appearing as a smear above 260 bp (21.0% of total fragments). The DSP-S cir-nDNA profile of HI had a major population between 85 and 260 bp (89.2% of total fragments), peaking at 166 bp (2.5% of total fragments); a minor population was also detectable between 261 and 420 bp (10.5% of total fragments), with no fragment detected above 420 bp. Periodic sub-peaks every 10 bp from 102 to 152 bp were detectable with cir-nDNA in contrast to cir-mtDNA. Cir-mtDNA and cir-nDNA SSP-S fragments size profile of HI ranged respectively from 50 to 475 nt and 40 to 400 nt (Fig. 2C, D). The cir-mtDNA SSP-S profile of HI had a major monomodal population between 50 and 150 nt, peaking at 80 nt, while there was a slowly decreasing population smear from 151 to 475 nt (25.2% of total fragments). Cir-nDNA SSP-S mean size profile had a major population between 45 and 260 nt (96.3% of total fragments), peaking at 166 nt, corresponding to nearly 2.0% of total fragments. The number of fragments plateaued between 70 and 120 bp at nearly 0.4% of total fragments. A very small population was observed between 261 and 400 nt (3.4% of total fragments). Periodic sub-peaks every 10 nt from 53 to 144 nt only existed with cir-nDNA.

**Figure 2:**
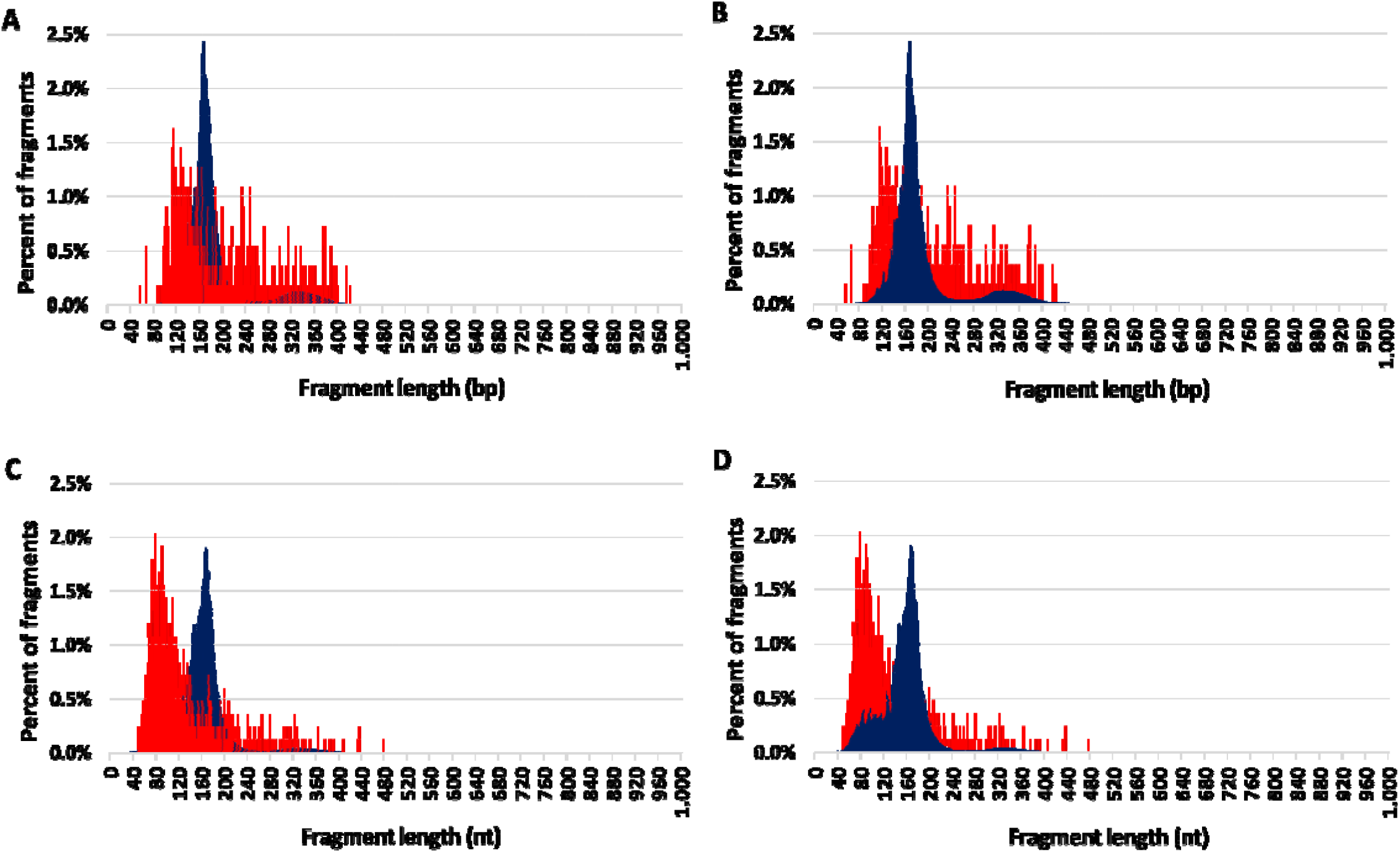
Comparison of the mean size profiles of circulating mitochondrial DNA (cir-mtDNA) and circulating nuclear DNA (cir-nDNA) from healthy individuals with alternate foreground. Low-pass WGS mean size profiles of cir-mtDNA (red) and cir-nDNA (dark blue) obtained with DSP-sequencing (A, B) and SSP-sequencing (C, D): cir-mtDNA in foreground and cir-nDNA in background (A, C); cir-mtDNA in background and cir-nDNA in foreground (B, D).

We observed a nearly 40-fold lower frequency of cir-mtDNA fragments < 90 bp with DSP-S (mean +/− SD = 0.9 +/− 1.2%) than with SSP-S (mean +/− SD = 33.5 +/− 4.9%) (Suppl. Table 2A and B). Additionally, the frequencies of fragments above 200 bp (nt) were homogenous among DSP-S (Suppl. table 2A) and SSP-S cir-mtDNA size profiles (Suppl. table 2B). However, the frequency of fragments above 200 bp (nt) was 3-fold higher with DSP-S (mean +/− SD = 38.1 +/− 4.2%) compared with SSP-S (mean +/− SD = 12.3 +/− 4.0%), meaning a higher number of fragments above 200 bp (nt) were obtained with DSP-S, compared with SSP-S. The ratio of the frequency below 90 bp over the frequency above 200 bp (Freq < 90) / (Freq > 200) was extremely homogenous among both DSP-S and SSP-S profiles. Nevertheless, this ratio was 134-fold lower with DSP-S (mean +/− SD = 0.02 +/− 0.03) than with SSP-S (mean +/− SD = 3.3 +/− 2.1).

### Physical examination of plasma prepared by various methods

The cirDNA-SPP as well as the PPw/oPA (as described in the Materials and Methods section) were partitioned to elucidate the fractions of the various cirDNA structural forms. Concentration values from plasma samples taken from five different healthy individuals, prepared under SPP or PPw/oPA, were directly compared to plasma prepared using only LS followed or not by F.

#### Effect of the use of HS and F on plasma prepared with the cirDNA-SPP

Supernatant cir-mtDNA concentrations following LS, LS+F, and LS+HS were 1.28, 0.02, 0.01 ng/mL, respectively. Thus, F and HS applied to the 1,200 g plasma supernatant resulted in a 98.7% and 99.4% decrease in cir-mtDNA level (Fig. 3A, B, Suppl. Table 3A). Values observed following LS were statistically different from both LS+F (p=0.0004), and LS+HS (p=0.0004), while no statistical difference existed between LS+F and LS+HS (Fig. 3A). Freezing led to a significant increase from 0.006 to 0.08 ng/mL (0.6% to 7.8 %), then a 12-fold increase of the cir-mtDNA amount in the 16,000 g supernatant, compared with LS+HS. As shown in Figure 3A, LS+freezing+HS values were statistically different from those of LS (p=0.0007) and LS+HS (p=0.01). Supernatant derived cir-nDNA concentrations following LS, LS+F, LS+HS and LS+freezing+HS were 5.28, 4.53, 7.25 and 9.27 ng/mL, respectively. No significant effect was observed on cir-nDNA level due to either filtration or 16,000 g centrifugation (Fig. 3A and B). Adding a freezing step between LS and HS increased by 28% the cir-nDNA concentration, which was not statistically significant (Fig. 3A, B). The proportions of cir-mtDNA mean concentrations among the total cirDNA mean concentrations were 19.5%, 0.3%, 0.1% and 0.9% following LS, LS+F, LS+HS and LS+freezing+HS, respectively. All concentration values are described in Supplementary Table 3A.

**Figure 3:**
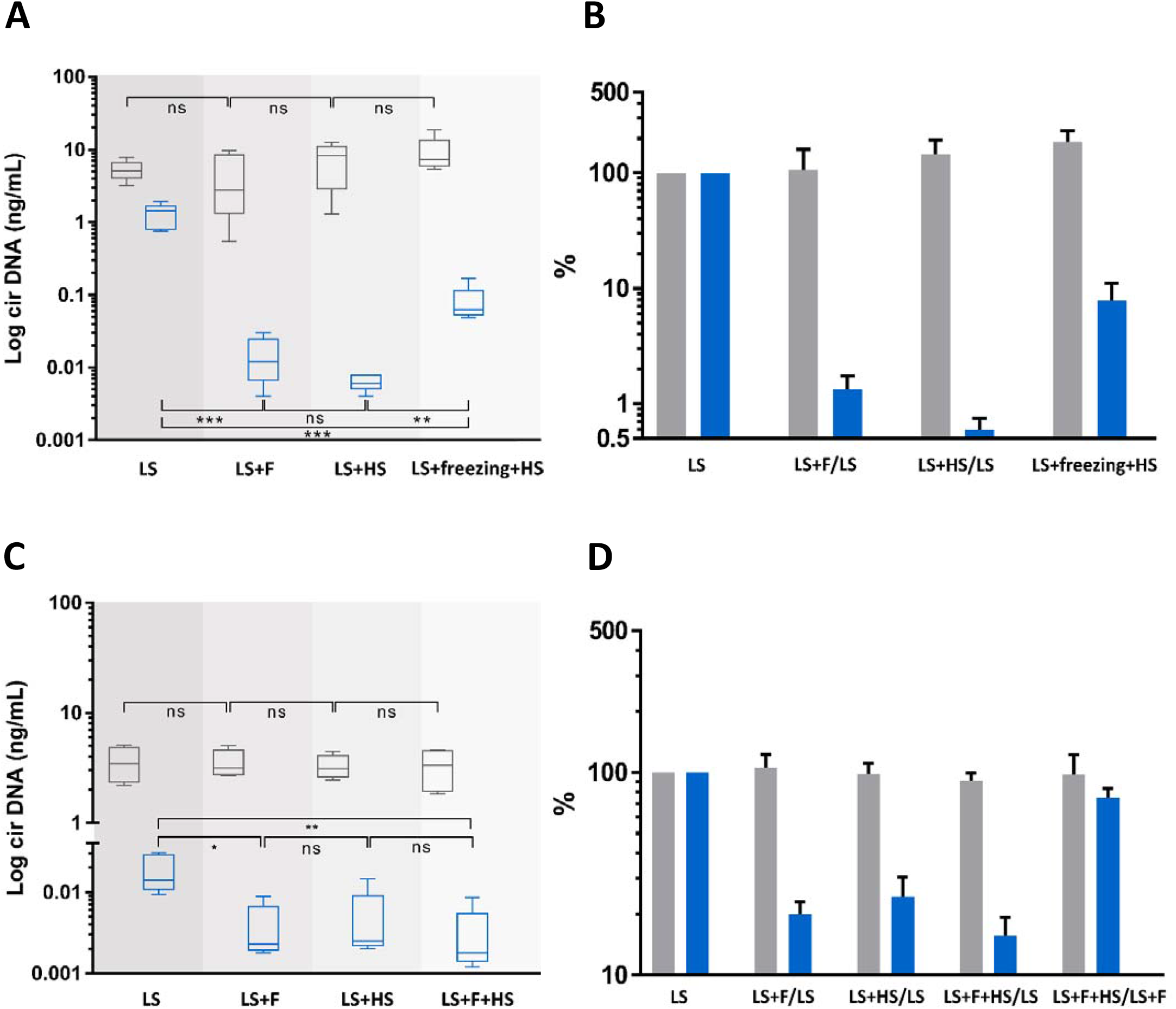
Comparison of the concentrations and variations of circulating nuclear (cir-nDNA, grey) and mitochondrial DNA (cir-mtDNA, blue) from 5 healthy individuals, depending on the plasma preparation protocol used and the physical procedure(s) applied subsequently to the resulting plasma samples. The preparation protocol used was either standard (SPP, A, B) or without platelet activation (PPw/oPA, C, D). The physical procedures applied were low speed centrifugation (LS), filtration (F), high speed centrifugation (HS) and freezing. Samples were either treated with LS only or with LS combined with F (LS+F), LS combined with HS (LS+HS) or LS combined with freezing and HS (LS+freezing+HS). The respective variation of the cirDNA concentration in each group was obtained comparing each combination with LS only, used as a reference. ns, non statistically significant; *p<0.05, **p<0.01; ***p< 0.001.

#### Comparison of the cirDNA-standard protocol and the protocol without platelet activation (PPw/oPA)

When using PPw/oPA, cir-mtDNA concentrations following LS, LS+F, LS+HS and LS+HS+F were 0.019, 0.004, 0.005, and 0.003 ng/mL, respectively. F and HS of the plasma following LS were statistically different to LS (p=0.012 and p=0.010, respectively) in terms of cir-mtDNA concentration, and resulted in a similar decrease (80 % and 75.7%, respectively) of cir-mtDNA levels (Fig. 3 C, D). By contrast, cir-nDNA quantification was not significantly altered by F or HS (Fig. 3C, D). In terms of amount, there were 67-fold less mtDNA detected in the supernatant of LS when using PPw/oPA as compared to SPP. In contrast, a much lower decrease of 1.5-fold was observed for cir-nDNA (Suppl. Table 3A, B). MtDNA detected in plasma following LS and HS showed a 199- and a 3.8-fold decrease of its concentration when using PPw/oPA and cirDNA-SPP, respectively. Note, a non-significant change was observed for cir-nDNA concentration (2.2-fold) after HS. In addition, F or HS altered the proportion of mtDNA by 64-fold and 128-fold, respectively, when using the SPP (Suppl. Table 3A). By comparison, it showed a nearly 5- and 4-fold decrease when using PPw/oPA (Suppl. Table 3B). The proportions of cir-mtDNA copies among the total cirDNA copies were 0.5, 0.1, 0.2 and 0.1% following LS, LS+F, LS+HS and LS+HS+F, respectively, using PPw/oPA. The proportion of cir-mtDNA to the total amount of cirDNA in the supernatant of the LS was much lower when PPw/oPA was used, compared to SPP (0.5% vs 19.5%). All concentration values are described in Supplementary Table 3A and B. Altogether, precluding platelet activation resulted in a decrease of about 377 million mitochondrial genome copies, standing for approximately a 98.5% reduction, as compared to SPP. About 99% and 76 to 80% of mitochondrial genome copies were lost following F or HS when comparing with SPP and PPw/oPA, respectively.

#### Estimation of extracellular vesicles associated cir-mtDNA proportion

While the plasma preparation without platelet activation provides highly qualitative information, its stringency, such as the numerous centrifugation steps, precludes accurate quantitative assessment of the cir-mtDNA. As a first step, we therefore used a Ficoll plasma preparation, and performed subsequent 16,000, 40,000 and 200,000 g centrifugations of the respective supernatant, which found about 3.7×10^6^, 9.2×10^5^ and 1.1×10^5^ copy number of mtDNA per mL, respectively. A statistical difference was only observed between the groups of plasma following the 16,000 g and the 40,000 g and between those following the 16,000 g and the 200,000 g centrifugations (p = 0.04, Fig. 4). From the total amount of mtDNA previously present in the 16,000 g supernatant, 24.1 % and 4.2 % of cir-mtDNA remained in the 40,000 g and 200,000 g supernatant, whereas 69.7% and 16.6% of cir-nDNA remained in the 40,000 g and 200,000 g supernatant (Suppl. Table 3C). The proportions of cir-mtDNA copies among the total cirDNA copies were 0.4, 0.1 and 0.1% following the 16,000 g, 40,000 g and 200,000 g centrifugation, respectively. All concentration values are described in Supplementary Table 3C.

**Figure 4:**
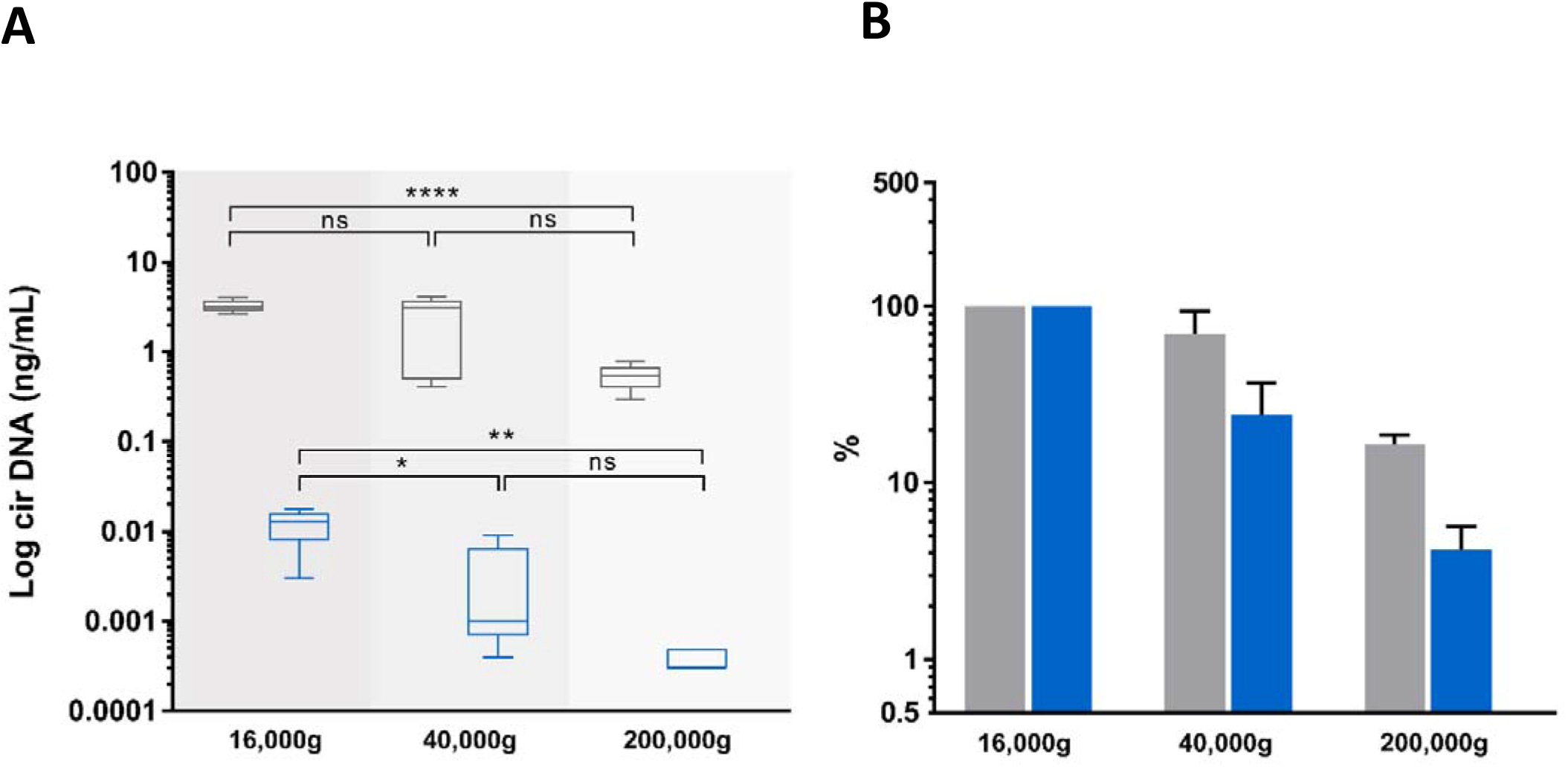
Effect of incremental centrifugation speed on plasma from 5 healthy individuals. Comparison of the concentrations (ng/mL), and variations (%) of circulating nuclear DNA (cir-nDNA, grey) and circulating mitochondrial DNA (cir-mtDNA, blue) in plasma as prepared by the protocol without platelet activation. cirDNA concentration was determined in the supernatant following the subsequent centrifugation steps of the respective supernatant at 16,000 g, 40,000 g and 200,000 g. ns, non statistically significant; *p<0.05, **p<0.01; ****p< 0.0001.

## DISCUSSION

Following the early discovery of cell-free DNA (cfDNA) and RNA in human serum by Mandel and Metais (1), a more recent study demonstrated the existence of another species of circulating nucleic acids, i.e. mtDNA (12). MtDNA’s structural as well as functional properties come from the peculiar mitochondria origin, which derives from the ancient symbiosis between free-living aerobic prokaryotes and eukaryotic cells. Note, other circulating nucleic acid species include viral, bacterial and fungal DNA or RNA. These whole types of circulating nucleic acids are now accurately characterized by high throughput sequencing (6,25,34,38,50,51). Despite previous efforts at designing methods, particularly Q-PCR based, for detecting mtDNA from blood cells (13,14,15,20), the standardization of cir-mtDNA quantification has proved to be a challenge, in particular concerning the issue of calibration. We recently designed an optimized method that enables the direct comparison of cir-nDNA and cir-mtDNA in human plasma (10). We also set out stringent pre-analytical conditions for this procedure (43). That previous study revealed that the plasma of healthy individuals contains approximately 50,000-fold more copies of the mitochondrial genome than that of the nuclear genome, while accounting for 10 to 25% of the total cirDNA mass content. This is consistent with the extremely high proportion of mtDNA molecules to the nuclear genome equivalent per cell (approximatively 1,000 to 20,000-fold), as well as the significantly lower size of the mitochondrial genome (16,569 bp) as compared with the nuclear genome (about 3×10^9^ bp).

Beyond quantitative observations, cir-nDNA analysis has benefited greatly from the understanding of its structural features acquired from their fragment size profile or fragmentomics. Works from Ellinger et al (52) and Diehl et al (53) revealed that fragments may be of a size lower than 180 bp; that was previously thought to be the lowest size (54), as it corresponds to that existing in a mononucleosome plus linker. Subsequently, we were the first to show that cirDNA is mostly highly fragmented, with a significant part of those fragments being of sizes down to 60 bp (44). Subsequent works using WGS analysis confirmed this in greater detail (25–27,50,51). In addition, WGS-assisted fragmentomics revealed nucleosome footprints, which indicated that detected short fragments (30 to 400 bp) derive from the nucleosome around which cirDNA is packed and relatively stabilized in the blood compartment (25,27,29). Furthermore, Shendure’s team (25) inferred tissue of origins from nucleosome occupancy as determined by WGS. We also described how cirDNA distribution in cancer patient plasma is mostly represented (67 to 80%) by mono-nucleosome associated cir-nDNA (29). In addition, Q-PCR assisted data on cirDNA distribution corresponded to WGS performed using SSP-S rather than DSP-S. This allowed the harmonization of data obtained from Q-PCR and WGS, since both SSP-S and Q-PCR use ssDNA as a template (28,29). A procedure combining LP-WGS and Q-PCR points to the possibility of identifying all kinds of nicks on fragments under 1,000 bp, and may also provide clues about potential cirDNA stabilizing structures in blood. Moreover, by adding a Q-PCR approach, we have given ourselves the possibility of learning more about fragments over 1,000 bp, and thus extending our comprehension of the size profile and the level of fragmentation of cir-mtDNA (28,29). As previously demonstrated (29), LP-WGS allows the inference of the presence of blunt or jagged dsDNA by comparing DSP-S and SSP-S. This process has also provided the following observations: (i), jagged dsDNA appeared by far the most present cirDNA structural forms; (ii), the lowest cirDNA fragment size is about 70 bp; and (iii), this might be due to the low hybridization forces of the lower-sized jagged fragments leading to their peeling off from histone/nucleosome particles (29). Altogether, our data suggest that the proportion of cir-nDNA inserted in mono-nucleosomes, di-nucleosomes, and chromatin of higher molecular size (<1000 bp) in healthy individuals ranges from 67 to 80%, from 9 to 12%, and from 8 to 21%, respectively (29).

Considering how useful this combined analytical approach had proved, we then applied it in an investigation of cir-mtDNA structural features. To this end, we used the cirDNA-SPP and a stringent clinically validated DNA extraction process, and followed our previously published guidelines (43). As we observed with cir-nDNA from LP-WGS, cir-mtDNA size profiles appeared homogenous and reproducible in the plasma of healthy individuals, whether detected by DSP-S or SSP-S; and in both cases, an overwhelming majority of fragments were less than 400 bp in size. This suggests that this population of short fragments is the result either of degradation by-products or of some means of cir-mtDNA protection. However, we found periodic distribution neither with DSP-S nor with SSP-S, while the DSP-S and the SSP-S mean size profiles differed significantly. With SSP-S, we detected a higher number of short fragments in the 50 – 120 nt interval, an earlier peak at 80 nt, and a faster decrease, as compared with DSP-S. DSP-S showed a higher proportion of fragments in the 120 – 200 bp interval, with a later and lower peak, around 120 bp, and a subsequent slower decrease. The significant differences between DSP-S and SSP-S may reveal a structural difference, since observation of a short population implies the protection of DNA strands by a stabilizing structure, as observed in cir-nDNA associated with nucleosomes. The clear differences existing between DSP-S and SSP-S profiles led us to believe that shorter ssDNA fragments appeared following SSP-S from “jagged” dsDNA, with at least one nick in one DNA strand still bound to the structure protecting the DNA from nuclease attack.

Only a few existing reports contributed to characterize cir-mtDNA structure. Chiu et al (55) initiated efforts towards this goal, and indirectly showed that cir-mtDNA may consist of both particle-associated and non-particle-associated forms of mtDNA in plasma. By paired-end sequencing analysis, a report (26) indicated that microbial and mtDNA are exposed to similar degradation processes, having observed that the fragmentation profiles of microbial and mtDNA in plasma were highly similar. Like Jiang et al (51), they also reported that cir-mtDNA size distribution is consistently shorter than cir-nDNA. Since mtDNA cannot be packed as nDNA with histones within nucleosome, both stated that cir-mtDNA is more susceptible to enzymatic degradation, given that its size profile reflects traces of wide range-sized mtDNA under dynamic degradation. Furthermore, Chandrananda et al found that SSP-S is more effective than DSP-S at recovering bacterial, viral and mitochondrial cirDNA (26); they also suggested that the fragmentation profiles of microbial and mtDNA in plasma are highly similar, indicating that they are exposed to similar degradation processes. These results may challenge the common paradigm, which considers fragments of cir-mtDNA below 400 bp as consisting only of degradation products (26). Burhnam et al later confirmed this statement (38).

Uncertainty as to the nature of the short cir-mtDNA fragments population nonetheless appears of minor importance, since its proportion to the total cirDNA fragments is very low: (i), the cir-mtDNA mean number corresponded to 0.006% and 0.012% of the total cirDNA fragments, when using DSP-S and SSP-S, respectively; (ii), there are nearly 50,000-fold more copies of the mitochondrial genome in plasma, which accounts for 10 to 25% of the total cirDNA mass content; and (iii), the DII as determined by Q-PCR showed that nearly all cir-mtDNA is over 310 pb. In contrast, cir-nDNA is highly fragmented with a DII showing a proportion of the number of detected fragments above 320 bp ranging from 10 to 20% (10,29). Since practically no cir-mtDNA fragments were detected between 310 bp and 1,000 bp, which is the practical upper limit of read-out in LP-WGS sizing using either DSP-S or SSP-S, we infer that most cir-mtDNA fragments are over 1,000 bp. Our data converge with that of several other studies. For instance, using a Q-PCR-based assay, Ellinger et al reported a plasma mtDNA integrity between 0.5 and 1.0 (56). Data obtained in this study confirmed our previous observation that the cir-mtDNA DII is always close to 1; this indicates that cir-mtDNA is not or very poorly fragmented (36). Note, we previously observed a much higher stability of cir-mtDNA compared to cir-nDNA extracted from serum-containing media of cultured cells (32).

The 3 main methods available for determining DNA fragment length are LP-WGS, Q-PCR, and Agilent capillary electrophoresis; however, these are limited in respect to (i) the practical upper limit of read-out around 1,000 bp (LP-WGS); (ii) the general low accuracy and precision (capillary electrophoresis); and (iii), the capacity to compare size fragments over 1,000 bp (Q-PCR). In order to decipher the major structural forms of the detected cir-mtDNA, we carried out a variety of experiments, based on physical examination and various plasma preparations. This led to a number of striking observations. First, we observed that, compared to the consecutive two-step centrifugation, the frozen storage step between LS and HS, as conventionally performed in the cirDNA-SPP, led to an increase by at least 10-fold of cir-mtDNA plasma concentration, whereas similar levels of cir-nDNA concentration were found. It should be noted that very recent work has reported similar observations (57). This means that freezing disrupts some structures in a way that leads to the release of cir-mtDNA but not cir-nDNA. Second, HS dramatically reduced the resulting plasma cir-mtDNA concentration by about 99 %, as compared to the 1,200 g plasma supernatant, whereas both supernatants revealed equivalent levels of cir-nDNA. This means that cir-mtDNA are mostly contained in or associated with structures whose densities correspond to that of cell organelles, membrane debris or apoptotic bodies. Third, filtration of the plasma supernatant obtained following a 1,200 g centrifugation resulted in a loss of nearly 99 % of detected cir-mtDNA, whereas no significant change was observed for cir-nDNA. This means that most of the cir-mtDNA are associated with structures whose size is over 0.22 μM. Note, our observations point to the existence of particles containing mtDNA in the circulation, as it has been previously indicated by plasma filtrates (55). In addition, our data agree with Arance-Criado’s observation (58), in healthy individuals, of a 98.7 % decrease of cir-mtDNA following the HS step using a protocol equivalent to the cirDNA-SPP. Taken together, these data suggest that nearly all detected cir-mtDNA derive from larger or denser biological structures, in contrast to cir-nDNA.

This suggestion should be linked to our recent discovery that blood contains circulating cell-free mitochondria (cir-exMT) (30). Data on genome integrity, specific protein immunoblotting, and fluorescent and electronic microscopy have all contributed to the evidence of the presence of this novel type of circulating blood component, which had previously gone unnoticed. Detection of the presence of intact cir-exMT in a normal physiological state was made possible by the Q-PCR’s strict specificity, allowing it to distinguish mtDNA from nDNA in plasma, as well as in cell culture media (30). Various works had previously reported cell-free DNA release from cells in culture medium (11,32,59). Cell culture studies enabled us to solve a particular question regarding the potential role of platelets in causing the presence of cir-exMT. This could originate either from passive platelet membrane disruption during the pre-analytical process, or from active release. Although no previous observation of mitochondria active release in normal physiological conditions from platelets existed prior to our work, there is a growing literature on the platelet release of cell-free mitochondria in specific pathological conditions (60,61). For instance, Boudreau et al previously demonstrated that platelets release mitochondria to serve as a substrate for the promotion of local inflammation (60). Puhm et al revealed the nature of an interferon I- and TNF-mediated inflammatory response by the release of extracellular mitochondria (exMT) and mitochondria embedded in microvesicles (MVs) by activated monocytes (61).

Hence, platelets are a source of cir-mtDNA as activated platelets can extracellularly release functional mitochondria. Various mediators regulate platelet activation (62) by controlling mitochondrial production of reactive oxygen species (ROS). Qin et al reported that platelets can release cir-mtDNA following their activation during cardiac surgery (63). In addition to platelet activation (60,62,63), the disruption of platelet membrane due to such physical factors as constraint forces or temperature may lead to mitochondria release. Increased detection of cir-mtDNA was found to correlate with platelet counts in either whole blood or PBMCs (18). Consequently, mature platelets which are devoid of nucleus but bear several mitochondrial genomes are confounders when quantifying cir-mtDNA from plasma. This is barely addressed in the literature concerning cir-mtDNA. Zhao et al reported that morphologically intact cir-exMT account for a significant majority of MVs in cases of traumatic brain injury (64). In addition, they revealed that exMT are bound to platelets and have a physiological role in the trauma-induced brain coagulopathy and inflammation (64).

In order to fully confirm cir-mtDNA associated structures, we performed a physical examination similar to the one previously used to demonstrate the existence of circulating cell-free mitochondria (30); this involved the inspection of plasma isolated without platelet activation. Use of F and HS resulted in an 80 % and 76% decrease of detected cir-mtDNA, respectively; this contrasted with cir-nDNA, whose amount remained only slightly affected by both procedures. In plasma prepared using the cirDNA-SPP, 0.22 μM filtration and 16,000 g centrifugation similarly decreased cir-mtDNA concentration, which suggests that both plasma preparations eliminate similar mtDNA containing particles. However, the percentage decrease was higher when using the cirDNA-SPP, highlighting the presence of a higher proportion of contaminating platelets or exMT deriving from platelets activation. F or HS altered the proportion of mtDNA by 64-fold and 128-fold, respectively, when the SPP was used, whereas that proportion showed a 5- and 4-fold decrease when using PPw/oPA. Around 3.7 ×10^8^ mtDNA copies/mL in SPP appeared in addition after LS as compared to PPw/oPA (Supp. Table 3). Filtration and sedimentation following the 16,000 g centrifugation reduced from 4.5×10^6^ and 4.1×10^6^ (mean, 4.3×10^6^) the cir-mtDNA copy number when using PPw/oPA. SPP enriched the amount of exMT of 92-fold, compared to PPw/oPA and proportion of exMT were 99.5% and 73.2% following SPP and PPw/oPA, respectively (Supp. Table 3D). Based on the assumption that there are 2 to 10 mitochondrial genome copies per mitochondria, our data from PPw/oPA consequently indicates that about 0.43 to 2.1 million/mL of exMT could circulate in blood of healthy subjects as discovered by Al Amir Dache (30).

Aside from cir-exMT origin, plasma-derived mtDNA could originate from other structures: remaining cells, in association with cell debris or membranes, mitochondria, or extracellular vesicles (EVs). There are three main EVs subtypes: MVs, exosomes, and apoptotic bodies. They may be differentiated based upon their biogenesis, release pathways, content, and function (65), and may show significant differences in size. MVs, exosomes and apoptotic bodies typically range from 100 nm up to 1 μm, <200 nm and >1μm in diameter, respectively (65). Differential centrifugations allow their isolation: apoptotic bodies at a g-force of approximately 2,000 g; MVs at 10,000-30,000 g; and exosomes by ultra-centrifugation at 100,000-200,000 g. Note, MVs can be differentiated as large (LMVs) and small (SMVs) (65). We may assume that EVs could contain either exMT or fragmented mtDNA genome.

Using sequencing analysis, Newell et al demonstrated the presence of only intact full mitochondrial genomes in the plasma cirDNA fraction (66), confirming our previous observation (30). They speculate that mtDNA could be protected from degradation by circulating nucleases due to either EVs encapsulation or the circular nature of mtDNA potentially delaying its degradation (66). This study does not account for mtDNA platelet origin, however, as they used an SPP equivalent process. Based upon our observation that the proportion of cir-mtDNA of size below 1000 bp is very weak, we infer that fragmented mtDNA between 1,000 bp and 16,000 bp, the approximate mitochondria full length genome, is barely associated with EVs (<0.5%). In contrast, our data based on the PPw/oPA revealed that 1.7% of the cir-mtDNA is associated with exosomes, and 18.4% associated with microvesicles or small/dead exMT (Table 2). Thus, when taking together fragmentomics and plasma fractionation data, we infer that a fraction, at least, of EVs contains mitochondrial full length circular DNA or mitochondria particles that could be internally or externally associated with EVs. Thus, we do not preclude the possibility that mitochondria particle free mtDNA may exist in blood circulation in association with EVs. However, its amount corresponds to a minor fraction of the total cir-mtDNA. In previous work (30), our transmission electronic microscopy (TEM) examinations showed no evidence of exMT encapsulated in or associated with bilayer phospholipidic vesicles or membranes. Moreover, our study combining LP-WGS and Q-PCR analysis showed that fragmented mtDNA of size below 1000 bp exist in extremely small quantities (<1%). Consequently, our data associated with others (30,55,66) show that mtDNA detected in plasma correspond quasi-exclusively to cir-exMT.

**Table 2:**
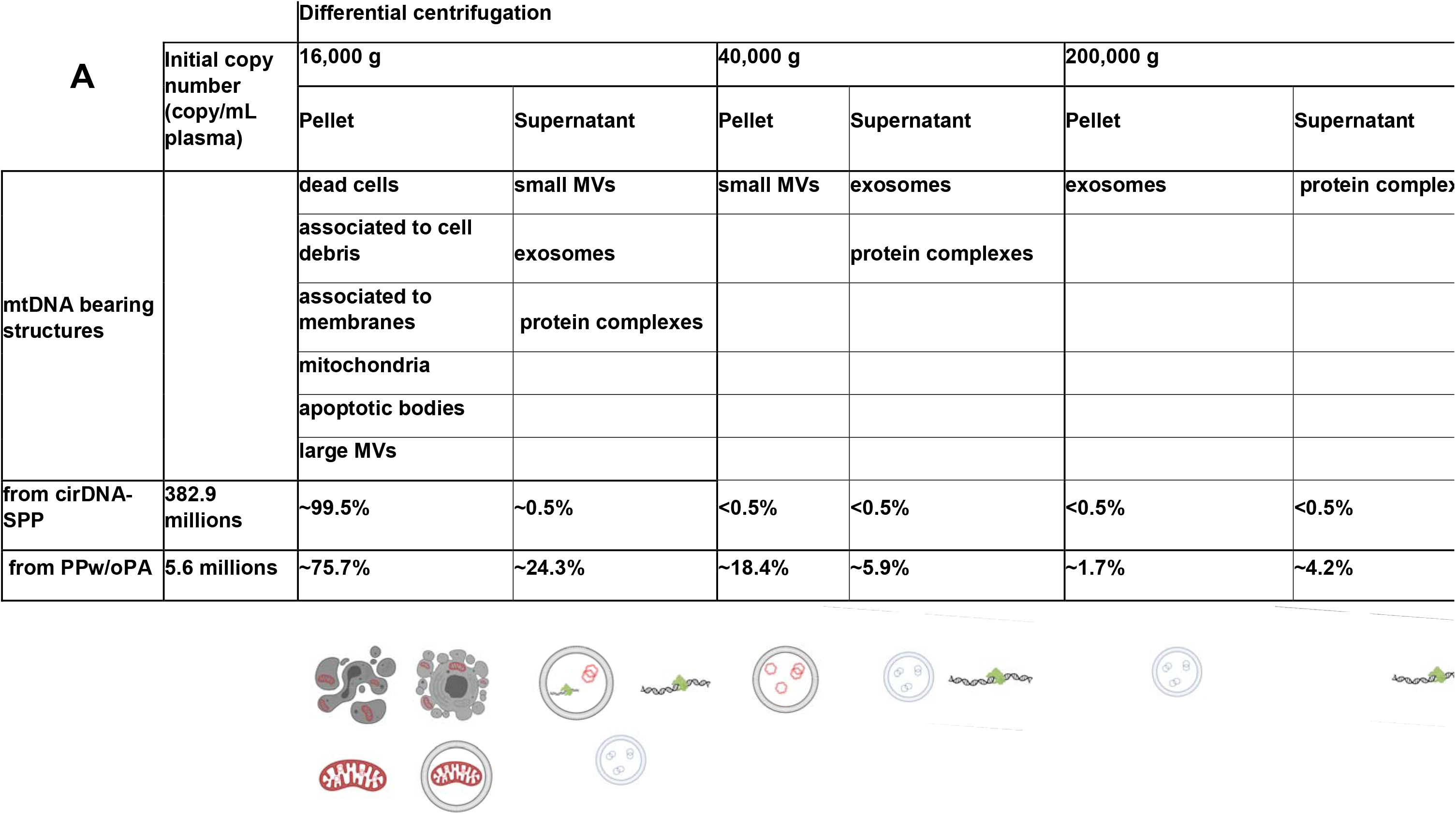
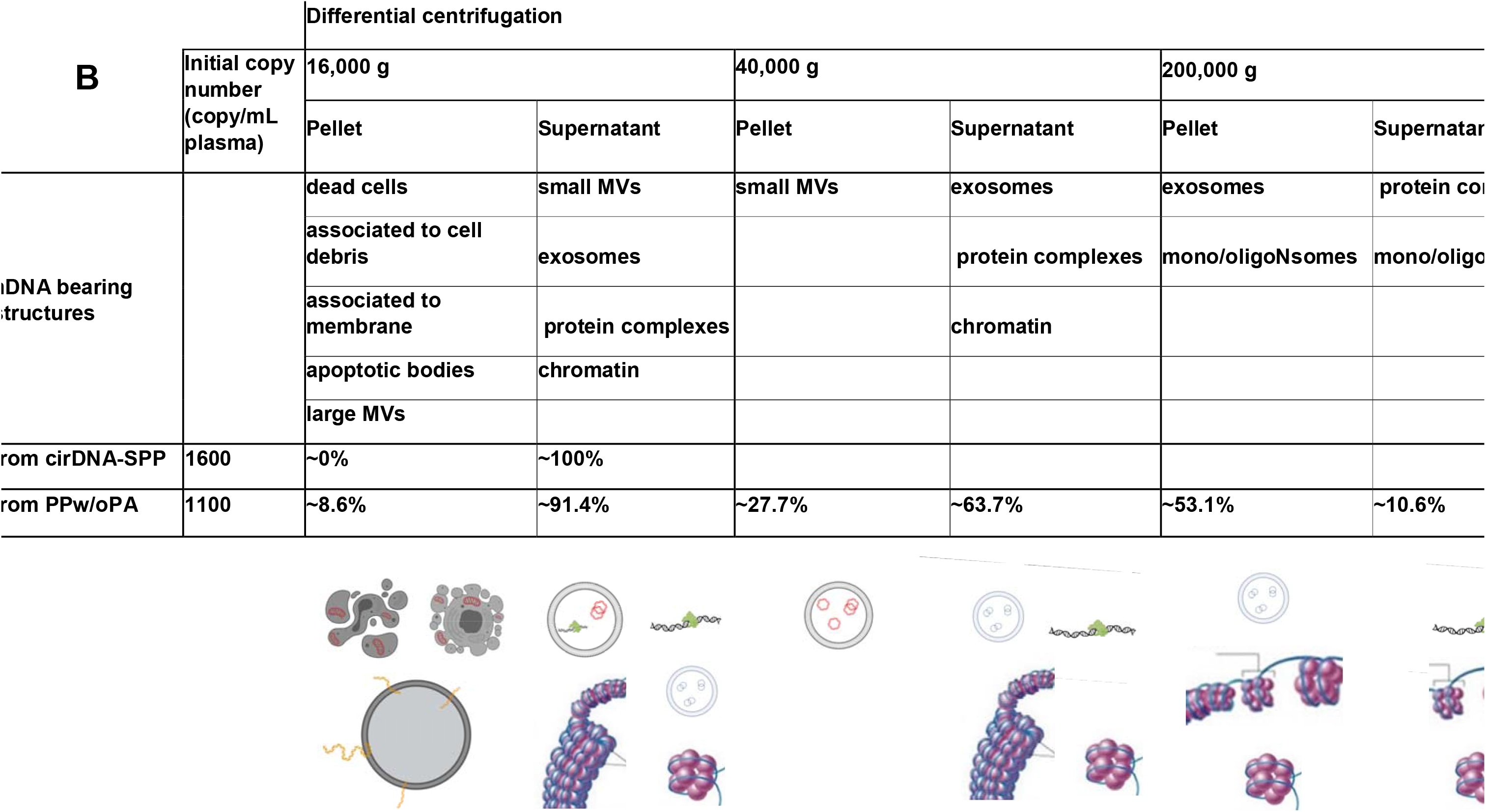
Suggested repartition of structures containing circulating DNA (cirDNA) in plasma. cirDNA content was measured in supernatant after successive centrifugations at different speeds, respectively 1,200 (initial copy number); 16,000; 40,000 and 200,000 g following either the cirDNA standard preparation protocol (cirDNA-SPP) or the preparation protocol without platelet activation (PPw/oPA). cirDNA content in the pellet was inferred from this of the supernatant at each step. A: mitochondrial cirDNA (cir-mtDNA); B: nuclear cirDNA (cir-nDNA); mtDNA, mitochondrial DNA; nDNA, nuclear DNA; MVs, microvesicles; oligoNsomes, oligonucleosomes.

The level of cir-mtDNA varied dramatically according to pre-analytical and analytical conditions. In particular, our data clearly shed light on the importance of the conditions of plasma preparation. The important points relevant to this issue are: (i), the plasma of the clinical routine preparation (involving only one LS) contains all cir-mtDNA structure types (exMT, EVs, protein complexes, highly degraded mtDNA); (ii); using a freezing step during the cirDNA-SPP appears to degrade exMT or mtDNA containing microparticles, which results in a higher plasma level following HS; (iii), while the plasma prepared without platelet activation contained much less cir-mtDNA (67-fold less), exMT nonetheless still represents the preponderant fraction of the total detected cir-mtDNA amount, compared to mtDNA containing EVs; and (iv), the paucity of the cir-mtDNA encapsulating microparticles/exosomes clearly confirms the need for specific isolation methods for their examination. The freezing step performed between the two centrifugation steps of the cirDNA-SPP led to a 12-fold increase of mtDNA content in the plasma following HS (Suppl. Table 3). We speculate that this fraction may consist principally of EVs-associated mtDNA, since EVs may have a much higher membrane disruption sensitivity to freezing than mitochondria (67), depending on which type of EVs is involved. Nevertheless, given the 7.8% of cir-mtDNA remaining in the supernatant following LS+freezing+HS, at least 92.2 % of mtDNA detected in the pellet following the cirDNA-SPP did not correspond to mtDNA-associated vesicles of low density, such as EVs encapsulating only fragmented mtDNA. The following confounding factors must also be taken into account: free mtDNA may be released by cell-free intact mitochondria undergoing degradation in blood; and platelets may actively release mitochondria under specific conditions (63,68). We are convinced that stringent experimental standardization is necessary in the use of cir-mtDNA as a biomarker or for the study of other biological processes. Besides applying the strict experimental conditions (42,43) we have used here, whenever plasma is used as a biological source (cirDNA-SPP), it is now common to employ a plasma isolation protocol with a double centrifugation step, as proposed by Chiu et al (55). Nonetheless, pre-analytical conditions such as the conditions (i.e. temperature, time delay…) of storage of the blood sample or of the supernatant following the first and second centrifugation step, and the use of cell stabilizing tubes, must be addressed as confounding factors. Those confounders are rarely taken into consideration, however, in the literature. Comparing or correlating observations from the literature as regards to the quantitative level of cir-mtDNA is therefore challenging, due to the lack of standardization in plasma preparation.

Consequently, our findings led to a profound revision of previous assumptions regarding the significance of the cir-mtDNA amounts detected in our and other previous works (10,34,51,55,59,69–71). First, it can no longer be considered valid to examine cir-mtDNA with WGS or to discuss cir-mtDNA fragmentation under 1000 bp when discussing the overall cir-mtDNA population, since it accounts for a very tiny part of the cir-mtDNA amount. Despite their low plasma content, these degradation products could nonetheless be relevant when considering differential enzymatic activity which may occur in certain physiological or pathological conditions, as suggested for cir-nDNA (29,72).

Despite the uncertainty about the true mtDNA level introduced by the confounding factors as revealed here, we remain extremely enthusiastic about cir-mtDNA’s potential diagnostic capacity for such conditions as inflammation diseases (73), neurological disorder (35,74), trauma (75), cardiovascular diseases (76), various cancers (40,58,77,78) and, as it was seen recently, SARS-Cov2 infection (79,80). The origins of cir-exMT, exMT in EVs, and vesicles encapsulating mtDNA still need to be distinguished, as the cir-mtDNA sources may depend on the biological observation, especially with respect to the nature of the plasma preparation protocols. Therefore, further efforts are needed to more fully characterize the features of these circulating biological structures, so that their diagnostic power as well as their biological function can be circumscribed.

Our study revealed that most of the authors, including ourselves, who studied this novel type of circulating biomarker previously had improperly assumed that they studied circulating fragmented mtDNA or mtDNA-associated EVs (10,33–35,40,58,63,75,77–79). We continue to believe, however, that highly fragmented cir-mtDNA degradation products as well as mtDNA containing EVs may have particular diagnostic relevance, despite their small proportions compared to that of cir-mtDNA deriving from exMT. Cir-mtDNA containing MVs have been isolated and intensively studied with regard to early diagnosis of various diseases, in particular of cancers or cardiovascular diseases (81,82). However, SPP appears to provide a mix of mtDNA-containing EVs and exMT. Conversely, cir-exMT may be isolated in the pellet following 16,000 g centrifugation, under a process precluding platelet activation (PPw/oPA), and may be numbered (30). Alternatively, cir-exMT may be estimated by assessing mtDNA content of the 2,500 g supernatant from the same PPw/oPA process. CirDNA-SPP was much more commonly employed in the literature, however, as was, to a lesser extent, MVs or exosomes isolation. Since exMT DNA was not suspected as the principal source of detected cir-mtDNA, it is reasonable to think that most biological observations implying mtDNA quantification rely on both cir-exMT, and the specific mitochondria constituents. Due to its microbial origin, cir-mtDNA as well as most of mitochondria constituents such as cytochrome C, ATP, succinate, N-formyl-peptides, mitochondrial transcription factor A (TFAM), or cardiolipin have been shown to constitute a powerful damage-associated molecular pattern (DAMP) (15,19,20,68). As previously demonstrated (60,61), exMT may also be regarded as released EVs representing critical intercellular mediators. Nevertheless, an uncontrolled release of cir-mtDNA in blood flow might be responsible for a pro-inflammatory state in a large spectrum of pathologies, including acute conditions such as systemic inflammatory response syndrome, acute liver failure, or chronic inflammatory diseases such as systemic lupus erythematosus, inflammatory bowel diseases, or cardiovascular diseases (19,20). Like microbial infections, these pathologies are associated with the release of nDNA and mtDNA from a ROS-dependent stimulation of neutrophils or eosinophils, leading to NETs formation (75). As a result, we speculate that cir-mtDNA, whether or not associated with mitochondria, may also have its origin in this innate immune response biological phenomena.

Since the observation that mitochondria are breaking through cellular boundaries, the characterization and quantification of cir-exMT may reveal mechanisms and functions of intercellular mitochondrial transfer. Recent reports highlighted the bioenergetic crosstalk between cells leading to procoagulant property, regulation of adipose tissue homeostasis, distressed cell rescue, as well as the therapeutic potential of mitochondrial transplantation (83–85). In particular, Brestoff et al (85) suggested that intercellular mitochondria transfer might have evolved as a homeostasis process implying immune cells in the regulation of local tissue microenvironment, and that targeting intercellular transfer might be considered as a future therapeutic strategy. Thus, cir-exMT appears as an exciting new biomarker in health and diseases.

We believe that certain controversies in previous literature arose from improper conclusions based on the confusion of cir-mtDNA and cir-nDNA, and the belief that at least part of their release derive from the same mechanisms. Consequently, we propose that future studies should systematically include a measurement of both entities, to circumscribe their respective physiological impact and diagnostic power. By comparison with cir-nDNA, the structural features of cir-mtDNA appear more complex and diverse, principally due to the release of exMT in interstitial milieu or blood circulation from various cell types, and due to the lack of stabilizing mitochondrial components which would enable protection from extracellular nuclease degradation. It is therefore necessary to better extend our knowledge of cir-mtDNA structure of origin, and to standardize the preparation of biological material, in order to fully determine and optimize the promise of cir-mtDNA in clinical and routine settings.

## DATA AVAILABILITY

The datasets used and/or analyzed during the current study are available from the corresponding author on reasonable request.

## SUPPLEMENTARY DATA

Supplementary Data are available at NAR online.

## ACKNOWLEDGEMENTS

The authors would like to thank:

- Nawel Yakoubi from MSD and Karine Saget and Vanessa Guillaumon from SIRIC Montpellier for their support.
- Charles Marcaillou and Steven Blanchard from Integragen for their help in obtaining sequencing datas.
- Marwin Edeas, Andreï Kudriatsev and Adil Sahla for their help during the preparation of this manuscript.
- Cormac Mc Carthy for his support and helpful comments on the manuscript
- Marc Ychou for his scientific support.

